# Investigating the links between trifluralin persistence in Western Australian soils and trifluralin resistance in resident annual ryegrass (*Lolium rigidum*) populations

**DOI:** 10.1101/2024.07.31.606087

**Authors:** Danica Goggin, Tim Boyes, Roberto Busi, Ken Flower

## Abstract

The pre-emergence herbicide trifluralin is widely used in the minimum-tillage cropping systems of Australia, with the result that resistance to trifluralin is increasing in the major weed of the region, annual ryegrass (*Lolium rigidum*). Repeated exposure to low herbicide rates is also known to result in the rapid evolution of resistance in weed populations. As trifluralin is highly volatile, readily photo-decomposed, metabolised by soil microbes and to bind strongly to soil organic matter, there are many factors that could result in weed populations receiving reduced (even sub-lethal) rates of the herbicide. To investigate whether trifluralin dissipation could play a role in the increasing levels of trifluralin resistance in annual ryegrass, resistance levels of populations from 18 Western Australian farms were compared with the dissipation rate of trifluralin applied to soil collected from these farms. Although there was no direct correlation between resistance level and trifluralin half-life, there were links between resistance and soil properties which suggest that higher rates of trifluralin dissipation could make a minor contribution to the development of resistance.

## Introduction

The major weed of the southern Australian cropping system, annual ryegrass (*Lolium rigidum*), is an obligate outcrossing species with a high level of genetic diversity and prolific seed production (Bajwa *et al*. 2021). These characteristics give it the ability to rapidly evolve resistance to the herbicides commonly used for its control (Owen *et al*. 2014; Broster *et al*. 2022), and, in response to the high levels of resistance occurring to selective in-crop herbicides such as the acetyl CoA carboxylase and acetolactate synthase inhibitors, growers are increasingly turning to pre-emergence herbicides (applied to the soil before crop sowing) to control their annual ryegrass populations (Boutsalis *et al*. 2014). One of the oldest pre-emergence herbicides is trifluralin, a dinitroaniline which inhibits binds to tubulin and prevents microtubule formation in the plant cell; this in turn causes severe inhibition of seedling coleoptile and radicle elongation, and prevents the seedling from establishing (Dostál and Libusová 2014). In response to repeated application of trifluralin over an extended period of time, resistance to this herbicide is becoming widespread in Australian annual ryegrass populations (Boutsalis *et al*. 2014), due to mutations in α-tubulin, which prevent the herbicide binding properly to its target site, and/or to enhanced detoxification of trifluralin by cytochrome P450 monooxygenases (Chen *et al*. 2021).

Trifluralin is a highly volatile compound which is usually incorporated into the soil upon application. Various studies on trifluralin dissipation from a range of soil types have concluded that volatilisation is the major pathway of trifluralin loss from soil (Mamy *et al*. 2005) and that the extent of volatilisation is negatively correlated with depth of soil incorporation (Savage and Barrentine 1969). High temperature and moisture conditions encourage volatilisation, whilst high soil organic matter slows dissipation (but also decreases bioavailability) due to the strong binding of trifluralin to organic compounds such as proteins and lipids (Weber 1990). Although it has been observed that biological degradation is not a major route of trifluralin dissipation overall (Mamy *et al*. 2005), it is known that fungi are the major soil microorganisms responsible for trifluralin degradation, and that degradation is more rapid under anaerobic than aerobic conditions (Weber 1990). Over a range of studies on trifluralin dissipation in the field, trifluralin half-life in the soil has been reported as ranging from 19 to 132 days (reviewed in Weber 1990).

Given that resistance to herbicides can develop rapidly when plants are exposed to sub-lethal rates (demonstrated in recurrent selection studies on both post- and pre-emergence herbicides, e.g. Neve and Powles 2005; Busi *et al*. 2012), it is of interest to determine if the rate of trifluralin dissipation from various soils is correlated with the resistance level of the resident annual ryegrass populations. For example, it is possible that in fields with a greater population of trifluralin-degrading microorganisms, annual ryegrass seedlings will be consistently receiving a sub-lethal dose of herbicide, accelerating the development of resistance. In the current study, dissipation of trifluralin applied to soils taken from a range of cropping fields in the grain belt of Western Australia was measured in a ‘common garden’ experiment, and compared with the trifluralin survival rate in potting mix of the populations collected from those fields.

## Materials and methods

### Trifluralin resistance data

The response of annual ryegrass populations from each soil collection location to the recommended rate of trifluralin was assessed as part of the UWA resistance testing service (see Busi *et al*. 2021 for a full description of the methods). All seeds were sown into the same sandy potting mix with a low organic carbon concentration (0.9%) and directly sprayed with trifluralin before being covered with fresh potting mix and watered. Seedling survival was recorded as a percentage of untreated controls for each population, with plants only considered as survivors if they had reached the 2-to 3-leaf stage at 28 d post-spray.

### Soil samples

The top 10 cm of soil (∼10 kg in total) was collected from eight farms in 2021 and ten farms in 2022, all located in the Western Australian grain belt and bounded by a rectangle of latitudes −30.71 to −31.57 and longitudes 116.6 to 118.1 (Fig. 1). Four replicate soil samples were collected from each farm in February of each year, allowed to air-dry if moist, and then sieved through a 2 mm mesh. The 2021 soils were kept at 4°C, with a subsample of each soil first sterilised by autoclaving at 121°C for 30 min. The 2022 soils were kept at room temperature and were not sterilised.

**Fig. 1.**
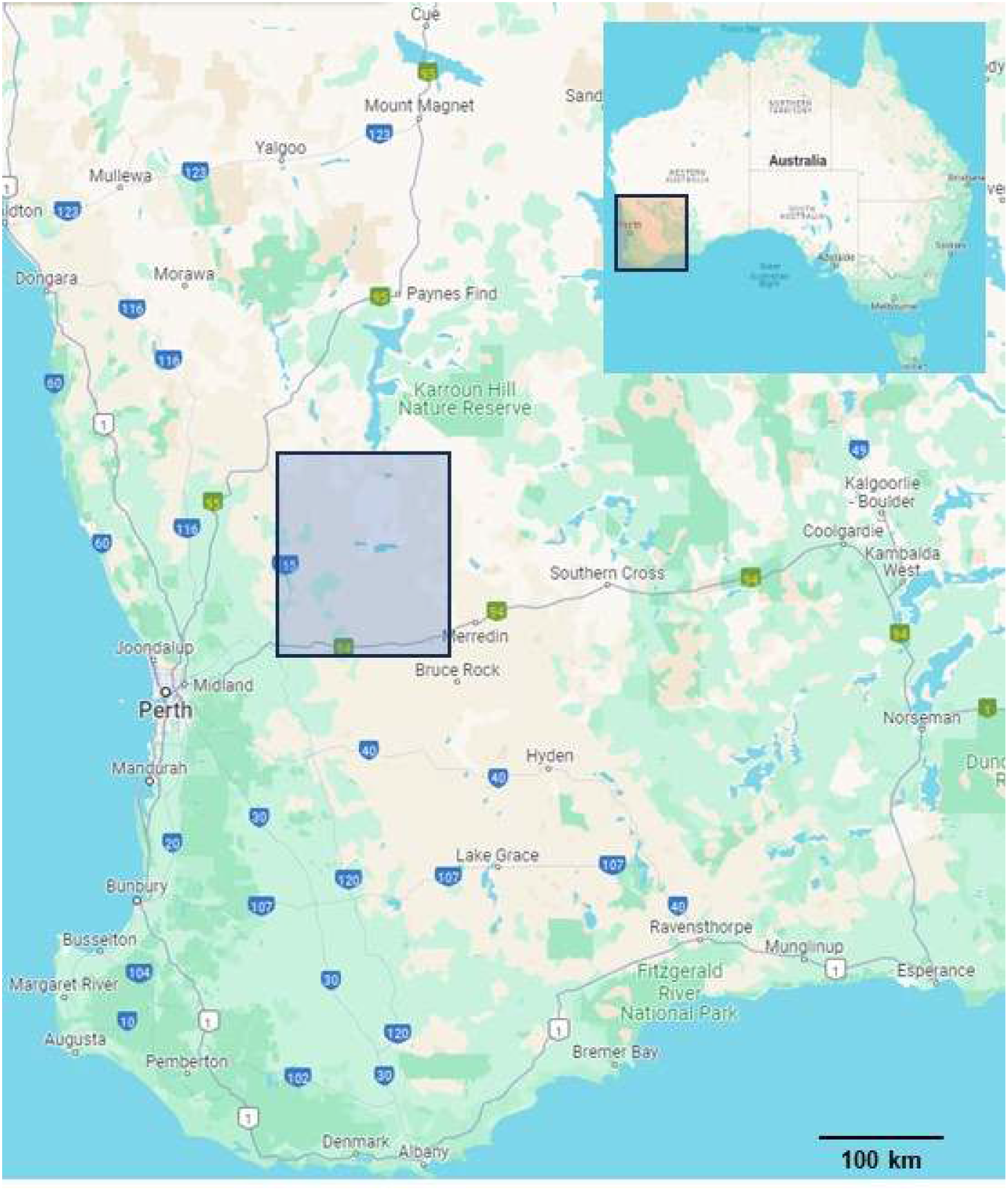
Map showing the area of soil sampling. The 18 farms in the Western Australian grain belt from which soils were sampled are bounded by the grey rectangle (inset: map of Australia showing the position of the main map in an orange rectangle). Maps generated by Google Maps.

### Analysis of soil properties

Soil particle analysis was performed according to Kettler *et al*. (2000). Briefly, 15 g of soil was agitated for 2 h in 3 vol of 3% (w/v) sodium hexametaphosphate and then passed through a 0.053 mm sieve. The sand retained by the sieve was oven-dried and weighed. The filtrate was stirred thoroughly, allowed to settle for 2 – 3 h, and the suspended clay fraction was decanted. The remaining silt was oven-dried and weighed, and the mass of the clay fraction deduced by subtracting the masses of the sand and silt fractions from the total starting mass.

Soil pH was measured in a 1:5 suspension of soil in 0.01 M CaCl_2_ after agitating for 60 min. Soil organic carbon was measured using the chromic acid wet oxidation method (Soon and Abboud 1991), where 0.5 g soil (dried for 4 d at 60°C) was ground to a very fine powder and added to 10 mL 166 mM K_2_Cr_2_O_7_. After the slow addition of 20 mL concentrated H_2_SO_4_, samples were incubated at 95°C for 45 min and the absorbance at 600 nm measured after diluting 300 μL of solution to 1 mL with water. Sucrose (0 – 48 mg), put through the same procedure, was used to construct a standard curve.

Soil microbial activity was assessed by measuring the hydrolysis of fluorescein diacetate (FDA) (El-Tarabily *et al*. 1996). Soil samples (5 g) suspended in 20 mL sterile 60 mM potassium phosphate buffer (pH 7.6) were incubated with 400 μL FDA, with FDA-free blanks included for each sample, at 25°C for 20 min. Reactions were stopped with 20 mL acetone, and the absorbance at 490 nm recorded. A fluorescein standard curve was made by hydrolysing 0 – 10 μg FDA in potassium phosphate buffer at 100°C for 30 min.

### Measurement of trifluralin dissipation over time

In 2021 and 2022, bioassays were employed to estimate trifluralin concentration in soil, using elongation growth of trifluralin-susceptible oat seedlings as the indicator (Cheam 1981). Three technical replicates of each soil sample (sterile and non-sterile for the 2021 samples) were used to fill the cells of 30-cell plastic trays, and watered to field capacity. These were then sprayed with formulated trifluralin (TriflurX, 480 g trifluralin L^-1^: Nufarm, Laverton North, Australia) at the recommended label rate of 960 g ha^-1^ and immediately covered with aluminium foil. Trays were kept outdoors during June (average maximum/minimum temperatures: 19/11°C), July (18/10°C) and August (19/10°C) in 2021 and May (22/12°C) and June in 2022. At various time points (0, 7, 14, 28 and 56 d after spraying in 2021; 0, 3, 7, 14 and 28 d in 2022), the foil was removed and five oat seeds per cell sown on the soil surface. Seeds were covered with 5 mm of the appropriate soil sample (untreated) and watered well. At 14 d after sowing, the radicle and coleoptile lengths of the oat seedlings were recorded. A standard curve was produced in each year by treating oat seeds on potting mix with 0, 9.6, 19.2, 96, 192, 480 or 960 g ha^-1^ trifluralin, with three replicates for each rate.

In 2024, trifluralin dissipation in all soil samples (non-sterile only) was re-analysed using direct measurement of trifluralin extracted from the soil. Moist soils were used to fill 100-cell plastic trays (one technical replicate per soil sample) which were then sprayed with 4800 g ha^-1^ trifluralin and immediately covered with aluminium foil. Trays were kept outdoors during April (25/15°C), May and June, 2024. At each time point (0, 3, 7, 21 and 63 d after spraying), the top 5 mm of soil was removed and allowed to air-dry for 24 h. Trifluralin was extracted according to Johnstone *et al*. (1998), by moistening 1 g of dry soil with 800 μL water and then incubating in 2.2 mL methanol for 60 min with gentle agitation. After allowing the soil to settle, a 1.5 mL aliquot was added to 6 mL water plus 1 mL saturated NaCl, and this was partitioned against 3 mL hexane. The organic phase (2.5 mL) was collected and the hexane evaporated under a stream of air. The residue was resuspended in 300 μL methanol and the absorbance at 426 nm (Escrig-Tena *et al*. 1998) recorded. Untreated samples of each soil were passed through the same extraction process and the A_426_ of the blank samples subtracted from that of the treated samples.

### Data analysis

Oat seedling radicle and coleoptile lengths were used to make a linear standard curve of tissue length as a percentage of the untreated control vs. log (trifluralin g ha^-1^). The inhibition of radicle elongation by trifluralin was so severe that only the coleoptile lengths could be used to calculate soil trifluralin concentration from the bioassay data.

Soil trifluralin concentrations derived from both the bioassays and direct measurement were used to calculate the half-life of the herbicide in each soil, based on the assumption that dissipation followed first-order kinetics (Mamy *et al*. 2005; Triantafyllidis *et al*. 2010) according to the equation:

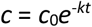

where *c* is the trifluralin concentration at time *t, c*_0_ is the concentration at *t* = 0 and *k* is the first-order rate constant. The trifluralin half-life was calculated by plotting ln(*c*/*c*_0_) vs. *t* and solving for *t* when *c*/*c*_0_ is 0.5 (Lehmann *et al*. 1993). The rate constant *k* is the slope of the linear regression line (Lehmann *et al*. 1993).

Correlations between parameters such as trifluralin half-life, soil properties and trifluralin resistance level were calculated using weighted least-squares regression in R (R Core Team 2023).

## Results

### Trifluralin resistance data and soil properties

Survival of annual ryegrass populations following treatment with the recommended rate of trifluralin ranged from 0 to 42%, with four farms having populations classed as susceptible (0 – 5% survival), nine farms having populations with developing resistance (6 – 19% survival), and the remaining five farms having populations classed as resistant (≥20% survival) (Busi *et al*. 2021) (Table 1). Soil organic carbon ranged from 1.3 – 2.8%, microbial activity from 3.3 – 9.8 μg FDA hydrolysed g^-1^ soil, pH from 5.0 to 5.9, and sand content from 18% to 81% (Table 1). Autoclaving of the 2021 soils resulted in microbial activity decreasing to 0 in all cases (data not shown).

**Table 1.** Trifluralin resistance levels in annual ryegrass populations from 18 Western Australian farms and the physical properties of the soils from the fields where the annual ryegrass was collected. Values are means ± SE (n = 4). Soil texture was classified according to the USDA soil texture triangle.

| Farm no. | Year | Ryegrass trifluralin survival (%) | Organic C (mg g<sup>-1</sup> dry soil) | FDA hydrolysis (μg g<sup>-1</sup> dry soil) | pH (measured in CaCl<sub>2</sub>) | Sand (%) | Silt (%) | Clay (%) | Soil texture |
| --- | --- | --- | --- | --- | --- | --- | --- | --- | --- |
| 1 | 2021 | 38 | 1.7 ± 0.1 | 7.5 ± 0.7 | 5.0 ± 0.1 | 51 ± 4 | 42 ± 4 | 7 ± 0 | sandy loam |
| 2 | 2021 | 42 | 1.7 ± 0.1 | 8.6 ± 2.6 | 5.3 ± 0.0 | 36 ± 2 | 54 ± 3 | 10 ± 1 | silt loam |
| 3 | 2021 | 2 | 1.8 ± 0.2 | 3.6 ± 2.1 | 5.4 ± 0.1 | 18 ± 3 | 71 ± 2 | 11 ± 1 | silt loam |
| 4 | 2021 | 28 | 1.8 ± 0.1 | 9.8 ± 0.2 | 5.2 ± 0.1 | 49 ± 6 | 43 ± 6 | 8 ± 1 | loam |
| 5 | 2021 | 42 | 2.0 ± 0.1 | 4.7 ± 0.3 | 5.3 ± 0.1 | 18 ± 1 | 76 ± 2 | 6 ± 1 | silt loam |
| 6 | 2021 | 15 | 2.0 ± 0.1 | 6.0 ± 0.7 | 5.5 ± 0.0 | 60 ± 4 | 32 ± 3 | 8 ± 1 | sandy loam |
| 7 | 2021 | 28 | 2.2 ± 0.0 | 5.0 ± 0.6 | 5.7 ± 0.0 | 56 ± 3 | 38 ± 4 | 5 ± 1 | sandy loam |
| 8 | 2021 | 0 | 2.5 ± 0.1 | 7.2 ± 0.6 | 5.7 ± 0.0 | 52 ± 2 | 41 ± 1 | 7 ± 1 | sandy loam |
| 9 | 2022 | 8 | 1.3 ± 0.2 | 3.6 ± 0.2 | 5.7 ± 0.2 | 81 ± 3 | 16 ± 3 | 3 ± 0 | loamy sand |
| 10 | 2022 | 6 | 2.4 ± 0.4 | 4.0 ± 0.6 | 5.7 ± 0.2 | 69 ± 2 | 27 ± 2 | 3 ± 0 | sandy loam |
| 11 | 2022 | 10 | 2.1 ± 0.2 | 4.1 ± 0.5 | 5.7 ± 0.2 | 62 ± 3 | 34 ± 3 | 4 ± 1 | sandy loam |
| 12 | 2022 | 6 | 2.2 ± 0.1 | 3.3 ± 0.2 | 5.5 ± 0.2 | 58 ± 3 | 38 ± 3 | 4 ± 0 | sandy loam |
| 13 | 2022 | 10 | 2.3 ± 0.1 | 6.2 ± 1.1 | 5.7 ± 0.2 | 63 ± 2 | 34 ± 2 | 2 ± 0 | sandy loam |
| 14 | 2022 | 12 | 2.6 ± 0.1 | 7.8 ± 0.4 | 5.4 ± 0.1 | 75 ± 4 | 23 ± 3 | 2 ± 0 | loamy sand |
| 15 | 2022 | 5 | 2.4 ± 0.3 | 6.1 ± 0.1 | 5.6 ± 0.1 | 75 ± 3 | 22 ± 3 | 2 ± 0 | loamy sand |
| 16 | 2022 | 6 | 2.1 ± 0.2 | 3.3 ± 0.1 | 5.7 ± 0.1 | 67 ± 5 | 30 ± 6 | 3 ± 0 | sandy loam |
| 17 | 2022 | 3 | 2.6 ± 0.2 | 4.6 ± 0.6 | 5.8 ± 0.1 | 61 ± 3 | 36 ± 3 | 3 ± 0 | sandy loam |
| 18 | 2022 | 14 | 2.8 ± 0.1 | 4.7 ± 0.7 | 5.9 ± 0.1 | 51 ± 7 | 47 ± 7 | 2 ± 0 | sandy loam |

### Trifluralin half-life in soils

In both years, the half-lives calculated from the bioassay data were short, ranging from 2 – 14 d, and were highly variable between replicates, with the regression lines for calculating first-order dissipation having an average r^2^ value of 0.898. This unreliability resulted in there being no significant differences between sterilised and non-sterilised soils, and few differences among farms (Fig. 2A). Therefore, trifluralin half-life was reassessed using direct measurement in the soil, which yielded an average of 55 d across all soils, a figure more in line with previous studies. The data for direct measurement appeared more reliable than that from the bioassays, with the average r^2^ of the regression lines being 0.945. However, there were still few significant differences among farms (Fig. 2B).

**Fig. 2.**
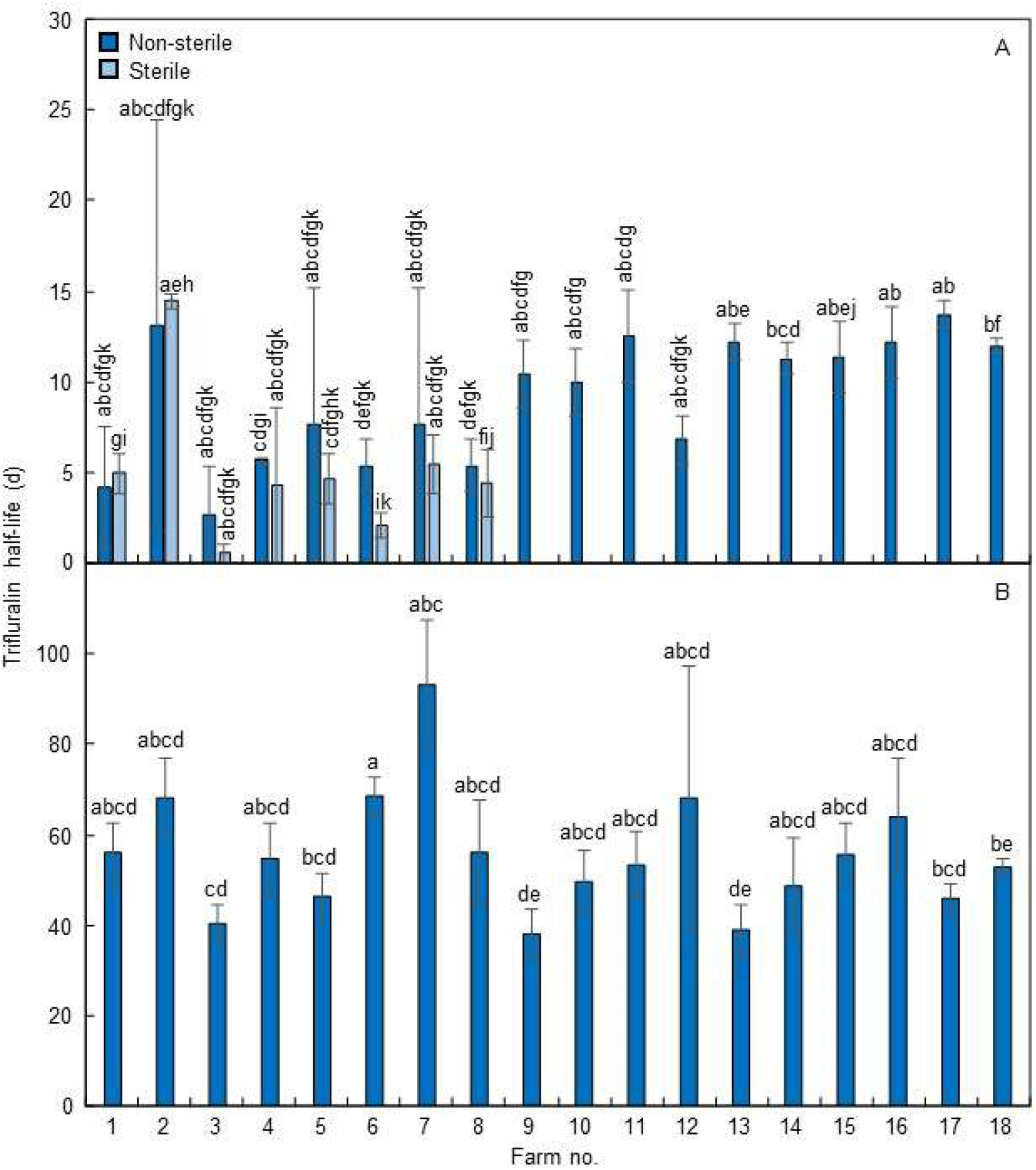
Half-life of trifluralin in soils collected from 18 farms in the Western Australian grain belt, measured using (A) a bioassay of oat coleoptile elongation in trifluralin-treated soil and (B) direct measurement of trifluralin extracted from the treated soil. Subsamples of eight of the soils were sterilised to determine if this affected trifluralin half-life (bioassay only). Values are means ± SE (n = 4); different letters above bars denote significant (*P* < 0.05) differences among means.

### Associations between trifluralin half-life, trifluralin resistance and soil properties

There was no correlation between the trifluralin survival of annual ryegrass populations from each farm and the half-life of trifluralin in the corresponding soils, using either the bioassay or direct measurement data for trifluralin half-life (Table 2). However, there were some significant correlations (*P* < 0.05) between trifluralin survival and the characteristics of the soil (microbial activity, pH, organic carbon and texture) from which the populations were collected (Table 2). There was a weak positive association between survival and microbial activity, and moderate negative associations between survival and soil organic carbon, sandiness and alkalinity (Table 2).

**Table 2.**
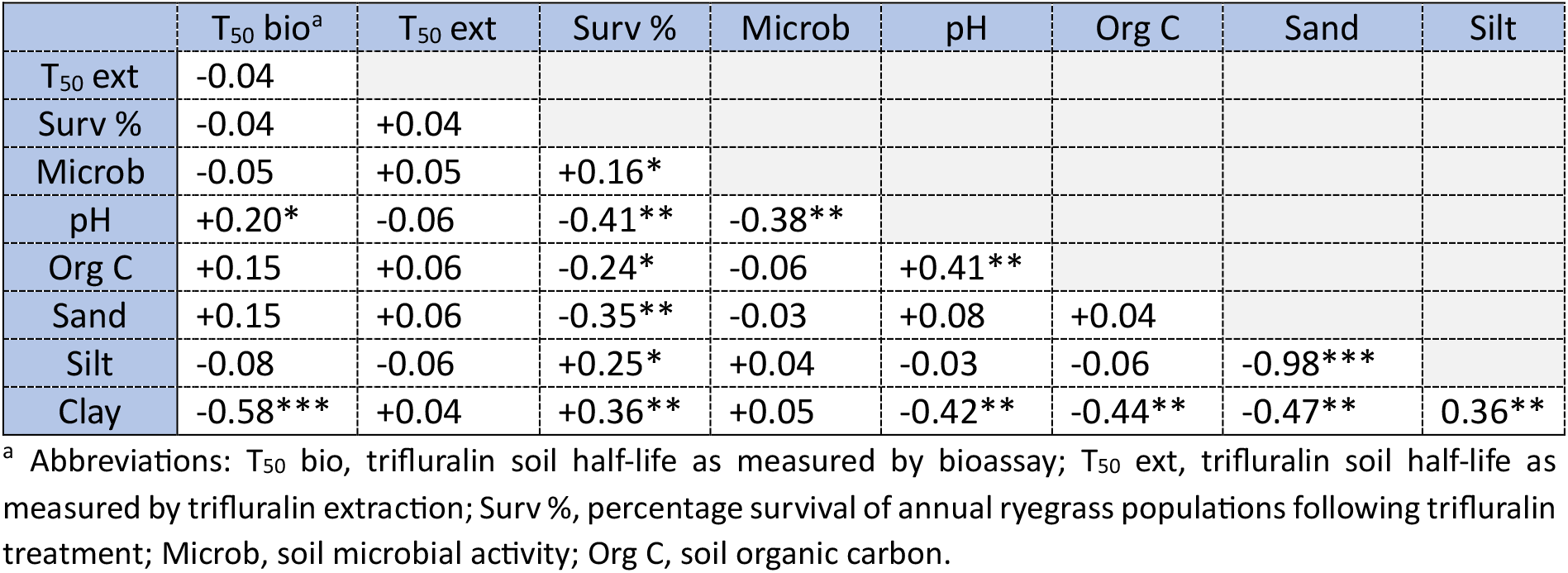
Correlation matrix for trifluralin soil half-life, survival of trifluralin-treated annual ryegrass populations and soil physical properties. Weighted least-squares regression was used to generate adjusted r^2^ values (positive and negative signs denote the direction of the correlation) between pairs of parameters. Asterisks indicate the significance level of each r^2^ value (none, *P* > 0.05; *, *P* < 0.05; **, *P* < 0.01; ***, *P* < 0.001).

## Discussion

Repeated exposure of highly adaptable weed populations to sub-lethal rates of herbicides can result in the rapid evolution of herbicide resistance due to the accumulation of resistance mechanisms in the population (Baucom and Busi 2019). Trifluralin, with its long history of use as a pre-emergence herbicide and its potential for rapid dissipation via volatilisation and photodecomposition and, to a lesser extent, microbial degradation (Savage and Barrentine 1969; Weber 1990), has become less effective on annual ryegrass populations in many southern Australian cropping fields (Broster *et al*. 2022). The aim of the current study was to determine if the level of trifluralin resistance in Western Australian annual ryegrass populations was associated with the persistence of trifluralin in the soil in which these populations grow, and if a higher rate of trifluralin dissipation could thus contribute to the faster evolution of resistance.

Using either a bioassay (elongation of oat seedlings sown into trifluralin-treated soil) or direct measurement of trifluralin extracted from the treated soils to calculate trifluralin half-life, there was no correlation between half-life and the ability of resident annual ryegrass populations to survive a trifluralin treatment. This could be at least partly due to the high variability among replicates. Sterilisation of some of the soils, although abolishing microbial activity, had no effect on trifluralin half-life. This suggests that, although microbial degradation of herbicides (e.g. trifluralin, profluralin, fluridone, prosulfocarb, carbetamide) has been known to decrease their persistence in the soil in previous studies (Camper *et al*. 1980; Freund *et al*. 1994; Rouchard *et al*. 1997; Hole *et al*. 2001), it is not the most important factor contributing to trifluralin half-life in the low-organic-matter Western Australian soils used in the current study. It must also be noted that the bioassay-based half-life calculations used for the sterilised vs. non-sterilised soils gave highly variable results which reduced the number of significant differences among samples.

Interestingly, there were significant correlations between the survival of annual ryegrass populations following trifluralin treatment and the physical properties of the soils from where the populations were collected, in spite of the fact that the resistance testing itself was performed in potting mix in a common garden experiment. It is therefore possible that the soil characteristics contributing to trifluralin performance (microbial activity, organic carbon, texture) do somewhat affect the amount of trifluralin reaching the germinating weed seedlings and thus have an influence on the enrichment of resistant individuals within a population. The (weak) positive correlation between trifluralin survival and microbial activity is in line with previous studies, but the negative correlation between survival and organic carbon was unexpected, given that soils with higher organic matter bind to trifluralin more strongly and thus make it less available to germinating seedlings (Peter and Weber 1985). The greater survival of trifluralin-treated annual ryegrass populations coming from soils of lower pH could potentially be explained by the fact that fungi are mainly responsible for microbial trifluralin degradation (Weber 1990), and these are the dominant microbes in low-pH soils (Camper *et al*. 1980; note also the significant correlation between soil pH and microbial activity in Table 2). Therefore, annual ryegrass populations coming from lower-pH soils could have been exposed to lower effective rates of trifluralin and thus gradually become enriched in trifluralin resistance mechanisms.

In conclusion, there are no convincing links between the trifluralin resistance level of Western Australian annual ryegrass populations and the persistence of trifluralin in the soils in which they occur, although there are indications that soil properties may have an association with the rate at which trifluralin resistance develops. As is often stated (e.g. Busi *et al*. 2021; Broster *et al*. 2022), proactive use of herbicide mixtures of multiple modes of action, in combination with non-chemical weed control techniques, can help to slow the evolution of resistance to individual herbicides and to control populations where resistance to a particular herbicide is already high. This will likely also help to slow the accumulation of soil microbes with the ability to rapidly metabolise pre-emergence herbicides such as trifluralin.

## Acknowledgements

This work was funded by the Grains Research and Development Corporation of Australia (project no. UWA2007-002RTX). We thank Professor Hugh Beckie for helpful suggestions in the early stages of the project.

## Competing interests

The authors have no competing interests to declare.

## Notes

### Competing Interest Statement

The authors have declared no competing interest.

